# *In vivo* imaging of mammary epithelial cell dynamics in response to lineage-biased Wnt/β-catenin activation

**DOI:** 10.1101/2021.06.22.449401

**Authors:** Bethan Lloyd-Lewis, Francesca Gobbo, Meghan Perkins, Guillaume Jacquemin, Marisa M Faraldo, Silvia Fre

**Affiliations:** Institut Curie, Laboratory of Genetics and Developmental Biology, PSL Research University, INSERM U934, CNRS UMR3215, F-75248 Paris Cedex 05, France; School of Cellular and Molecular Medicine, University of Bristol, Biomedical Sciences Building, Bristol, BS8 1TD, UK

**Keywords:** mammary gland development, lineage tracing, β-catenin, Wnt Signaling, intravital imaging, *in vivo* imaging

## Abstract

Real-time, *in vivo* imaging provides an essential window into the spatiotemporal cellular and molecular events contributing to tissue development and pathology. By coupling longitudinal intravital imaging with genetic lineage tracing, here we captured the earliest cellular events underlying the impact of active Wnt/β-catenin signaling on the organization and differentiation of the mammary epithelium. This enabled us to interrogate how Wnt/β-catenin regulates the dynamics of distinct subpopulations of mammary epithelial cells *in vivo* and in real time. We show that β-catenin stabilization, when targeted to either of the mammary luminal or basal epithelial lineages, invariably leads to cellular rearrangements that precipitate the formation of hyperplastic lesions that undergo squamous transdifferentiation. These results enhance our understanding of the earliest stages of hyperplastic lesion formation *in vivo*, and reveal that in mammary neoplastic development, β-catenin activation dictates a hair-follicle/epidermal differentiation program independently of the targeted cell of origin.

## Introduction

The Wnt/β-catenin pathway is a fundamental and highly conserved signaling cascade that regulates tissue morphogenesis and stem cell fate in several tissues. Wnt signal activation results in the accumulation and nuclear translocation of β-catenin, which acts as a co-transcriptional activator of TCF/LEF target genes important for cell proliferation, survival and differentiation (MacDonald et al., 2009; Nusse and Clevers, 2017). In the absence of a Wnt ligand, the pathway remains inactive, with cytoplasmic levels of β-catenin maintained low by continuous proteasomal degradation (Nusse and Clevers, 2017; Steinhart and Angers, 2018).

Wnt/β-catenin signaling plays a central role at all stages of mammary gland development (Incassati et al., 2010; Jardé and Dale, 2012; Yu et al., 2016), and aberrant pathway activity is also widely implicated in breast tumorigenesis. Wnt-1, the prototype member of the Wnt family, was originally identified as a site of integration by the mouse mammary tumor virus (MMTV) (Nusse and Varmus, 1982), providing the first link between Wnt signaling and breast cancer. Since, numerous studies have shown that dysregulated Wnt/β-catenin signaling leads to perturbed mammary gland development and tumorigenesis (reviewed in (van Schie and van Amerongen, 2020; Yu et al., 2016)). Targeted expression of Wnt-1 to the mammary luminal epithelium under the control of the MMTV long terminal repeats (LTR) results in a hyperbranched mammary phenotype and mammary adenocarcinomas (Li et al., 2003; Liu et al., 2004; Teissedre et al., 2009; Tsukamoto et al., 1988). Similarly, forced activation of Wnt signaling via MMTV-mediated expression of stabilized forms of β-catenin lacking N-terminal phosphorylation sequences (ΔN89- or ΔN90-β-catenin) results in precocious alveologenesis, and eventually adenocarcinomas (Imbert et al., 2001; Michaelson and Leder, 2001; Teissedre et al., 2009). Also, N-terminally truncated β-catenin (ΔN57) targeted to the mammary basal compartment using the keratin (K) 5 promoter induced basal-type hyperplasia in nulliparous aged females, in addition to squamous and invasive carcinomas in multiparous mice (Moumen et al., 2013; Teuliere et al., 2005).

In an alternative model, β-catenin stabilization can be achieved by Cre-mediated excision of loxP-flanked exon 3 of the endogenous *Catnb* gene (*Catnb*^+*lox(ex3)*^ mice) (Harada et al., 1999). Unlike models triggering exogenous β-catenin activation, *Catnb*+^*lox(ex3)*^ mice fail to develop mammary adenocarcinomas when β-catenin stabilization is induced in the luminal epithelium using whey acidic protein (WAP)-Cre (Pittius et al., 1988), or in all mammary epithelial cells with MMTV-Cre (Miyoshi et al., 2002b, 2002a). The impact of targeting mutant *Catnb*^+*lox(ex3)*^ specifically to the mammary basal epithelial compartment, and how this compares to phenotypes observed in luminal cells (Miyoshi et al., 2002b, 2002a), has yet to be investigated. As the molecular signatures and histopathological features of cancer cells do not necessarily reflect their presumptive cells of origin (Lim et al., 2009; Molyneux et al., 2010), studies focused on accurately dissecting the impact of the same oncogenic mutation in different cell types are warranted. Moreover, the early impact of sustained Wnt signaling on dynamic mammary epithelial cell behaviors and their neighboring wild-type cells, and how this affects the organization of the mammary gland during neoplastic transformation, is also largely unknown. Indeed, the rare nature of mutagenic events significantly hampers the *in situ* visualization of the earliest stages of cellular transformation. Thus, revealing the dynamic cellular mechanisms underlying this process promises to provide important insights into the critical steps leading to breast cancer initiation. To this end, we coupled genetic lineage tracing with longitudinal high-resolution intravital microscopy (IVM) to visualize the earliest changes in mammary luminal or basal epithelial cell dynamics in response to constitutive Wnt/β-catenin activation.

## Results

### Targeted stabilization of β-catenin to luminal mammary cells leads to hyperplastic lesions

The mammary ductal network is composed of two main epithelial lineages: basal and luminal cells, with the latter subdivided based on the expression or absence of the hormone receptors estrogen receptor-α (ERα) and Progesterone Receptor (PR). Notch signaling is a critical determinant of luminal cell fate (Bouras et al., 2008; Lilja et al., 2018), with Notch1 receptor expression restricted to ERα/PR-negative luminal progenitor cells in the postnatal mammary gland (Rodilla et al., 2015). Thus, to investigate the impact of constitutive Wnt/β-catenin activation on luminal progenitors, we crossed *Catnb*^+*lox(ex3)*^ mice (Harada et al., 1999) to Notch(N)1Cre^ERT2^;R26^mTmG^ mice (Rodilla et al., 2015) (Fig.1A, henceforth referred to as N1Cre/Tom). In these compound mice, the N1-Cre^ERT2^ line (Fre et al., 2011) is crossed to the double fluorescent reporter model Rosa26^mTmG^ (Muzumdar et al., 2007), enabling membrane-bound tdTomato expression to be switched to membrane-bound green fluorescent protein (GFP) in Notch1-expressing cells and their progeny upon low-dose Tamoxifen (TAM) administration (Fig. 1B, S1A) (Rodilla et al., 2015). While β-catenin was predominantly restricted to epithelial basolateral membranes in wild-type mice, tamoxifen administration in mutant N1-Cre^ERT2^;R26^mTmG^;β-catn^flox(Ex3)/+^ mice (henceforth referred to as N1Cre/β-cat) resulted in β-catenin accumulation in mammary luminal cells that correlated with GFP expression, which represented a robust indicator of mutant β-catenin status (Fig. 1B). As expected (Lilja et al., 2018; Rodilla et al., 2015), flow-cytometry analyses confirmed the localization of Notch1-derived GFP+ mammary epithelial cells to the luminal epithelial compartment (CD24^+^CD29^low^) in both models (Fig.1C, D Fig.S1B).

**Fig. 1.**
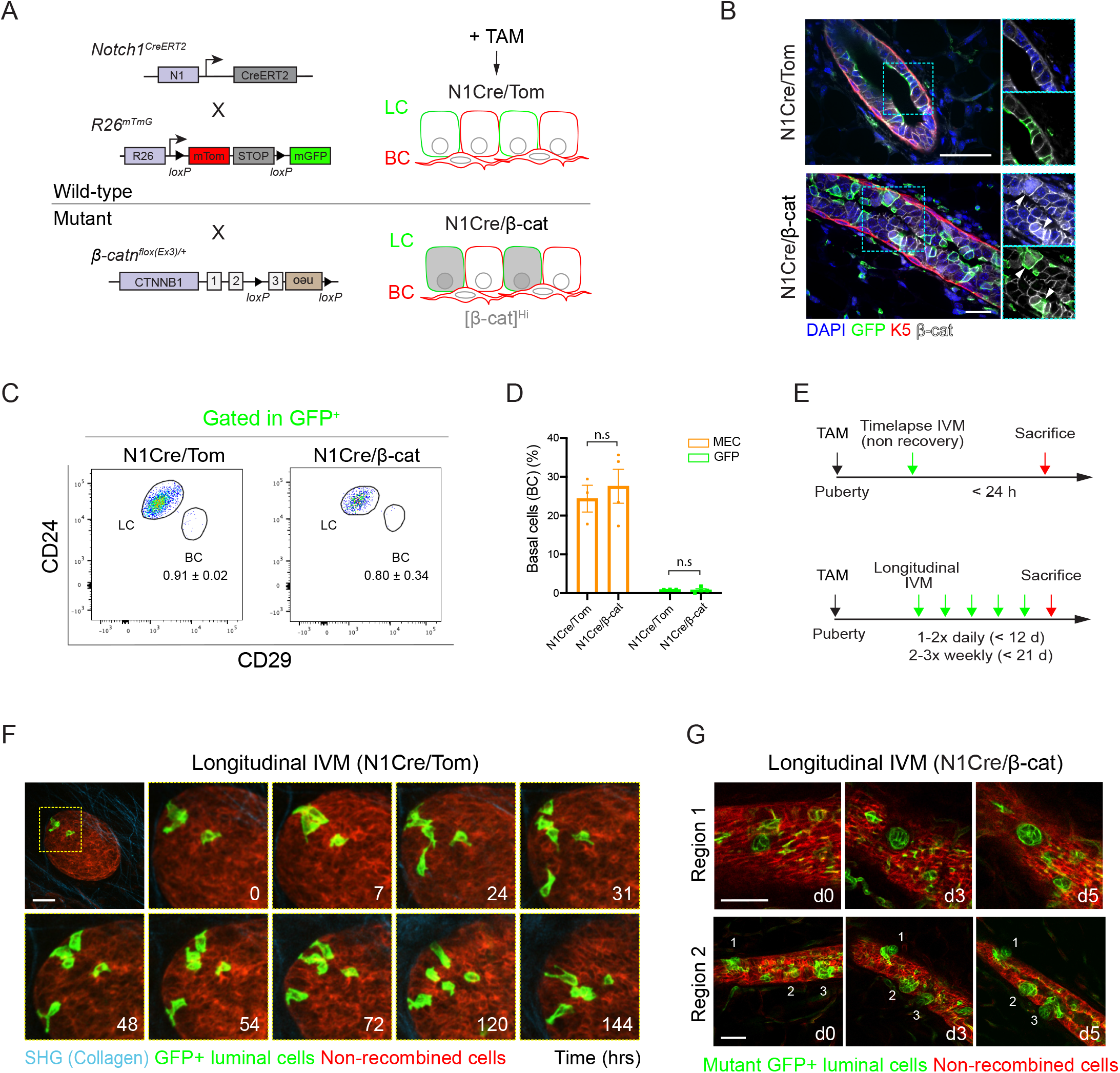
β-catenin activation in Notch1-expressing mammary luminal cells drives the development of mammary hyperplasia. **(A)** Schematic representation of the Notch1-Cre^ERT2^;R26^mTmG^ and Notch1-Cre^ERT2^;R26^mTmG^;β-catn^flox(Ex3)/+^ models used in this study (referred to as N1Cre/Tom and N1Cre/β-cat, respectively). All cells express membrane tdTomato fluorescence (red). Low-dose Tamoxifen (TAM) administration induces membrane GFP (green) labelling of sparse Notch 1-expressing luminal cells and their progeny in both models, and mutant β-catenin accumulation in N1Cre/β-cat mice (grey). **(B)** Representative sections of mammary gland tissues showing β-catenin cytoplasmic/nuclear accumulation in luminal cells in the mammary epithelium of N1Cre/β-cat mice 7 days after TAM induction, which coincided with membrane GFP expression (arrowheads). K5 is shown in red, anti-β-catenin staining in white and DAPI labels nuclei in blue. Scale bars: 20μm. **(C)** Representative FACS dot plots of luminal CD24^+^CD29^lo^ (LC) and basal CD24^+^CD29^hi^ (BC) mammary cells gated within the GFP+ mammary epithelial cell (MEC) population in pubertal N1Cre/Tom and N1Cre/β-cat mice 48-72 h after tamoxifen administration. GFP+ MECs are restricted to the luminal CD24^+^CD29^low^ compartment in both models. Average values are shown ± SEM. **(D)** Quantification of GFP+ CD24^+^CD29^hi^ cells (green) as compared to the proportion of total BCs within MECs (orange) in N1Cre/Tom or N1Cre/β-cat 48-72 h after tamoxifen administration. Graph shows mean ± SEM. No statistical differences (n.s) were observed between transgenic lines (p > 0.05, Welch’s t-test, n=3-4 mice per group). **(E)** Schematic representation of experimental timelines for time-lapse or longitudinal intravital imaging of mammary glands in wildtype and mutant β-catenin models. **(F)** IVM images of a mammary end bud structure in a pubertal N1Cre/Tom mouse showing recombined GFP+ (green) mammary luminal epithelial cellular rearrangements over 6 days (144 h). Red; non-recombined membrane tdTomato-expressing mammary epithelial cells, cyan; collagen (SHG). n = 4 mice. Related to Supplemental Movie 1. Scale bars: 50μm. **(G)** IVM images of a mammary duct in a N1Cre/β-cat mouse showing the development of hyperplastic luminal GFP+ (green) lesions over time, n = 3 mice. Day (d)0 represents 16 days after Tamoxifen induction. Red; non-recombined membrane tdTomato-expressing mammary epithelial cells. Scale bars: 50μm.

To visualize cellular dynamics *in vivo* and *in situ* during mammary ductal development, we surgically implanted an imaging window (MIW) (Jacquemin et al., 2021; Zomer et al., 2013) over the abdominal (4^th^) mammary gland of 5-6-week-old transgenic mice, 24-72 h after tamoxifen administration (Fig. 1E). This approach enabled the high-resolution 4-dimensional (x-, y-, z- t-) intravital imaging of the mammary epithelium over time in physiological conditions. Consistent with recent short-term IVM studies using neutral labelling strategies (Corominas-Murtra et al., 2020; Messal et al., 2021; Scheele et al., 2017), longitudinal IVM in pubertal mammary glands of N1Cre/Tom mice revealed the cellular rearrangements of terminal end bud (TEB)-resident luminal cells (Fig. 1F, S2A, Movie S1), validating our *in vivo* imaging conditions. Interestingly, Notch1-derived GFP+ luminal progenitors in stratified TEBs intermittently, but regularly, extended cellular protrusions to contact the basal epithelial layer over several days, exposing them to signals from the basement membrane (Fig. 1F, S2A, Movie S1), as previously observed in fixed tissues (Lafkas et al., 2013) and by time-lapse imaging of mammary organoids (Ewald et al., 2012). Notably, these cells were maintained at the distal tips of TEBs, and not deposited in subtending ducts during branching morphogenesis. Notch1-derived luminal GFP+ cells in ductal structures were also observed to dynamically interact with the basal compartment by longitudinal intravital imaging spanning several days (Fig. S2B). This serial imaging approach also revealed at an unprecedented resolution the dynamic process of mammary lumen formation *in vivo*, whereby fusion of several discrete lumina drives the establishment of the ductal network (Fig.S2C-D).

Our ability to visualize the cellular dynamics of individual mammary epithelial cells *in situ* over time by IVM provided a powerful platform with which to interrogate the early impact of sustained Wnt signaling on the dynamics of specific luminal or basal mammary cells. We therefore coupled the lineage tracing of β-catenin gain-of-function mice with intravital imaging of mammary epithelial cell dynamics. Although this low-throughput approach poses challenges for quantification, time-lapse imaging of mutant GFP+ mammary luminal cells in N1Cre/β-cat mice suggested their increased mobility compared to wild-type luminal cells (Fig. S3A, Movie S2-4). Mammary epithelial structures, however, appeared morphologically normal at this stage (Fig 1B, S3A). To visualize the long-term impact of Wnt/β-catenin activation on luminal cell dynamics, we performed longitudinal IVM in the mammary glands of pubertal N1Cre/β-cat mice for up to 3 weeks (Fig. 1E). This approach revealed the dynamic rearrangement of GFP+ mutant cells into compact, budlike clusters that expanded over time, rapidly encroaching neighboring basal layers (Fig. 1G, d0 = 16 days after TAM induction). Ring-like lesions with empty central regions were also observed, suggesting that internal cells within developing growths may undergo cell death (Fig. S3B, 19-21 days post induction). Collectively, these findings show that activated Wnt signaling promotes luminal epithelial cell rearrangements to drive the development of hyperplastic mammary lesions.

### Targeted stabilization of β-catenin to basal mammary cells also leads to abnormal epithelial budding and hyperplastic lesions

To induce β-catenin stabilization specifically in basal cells, we crossed the same *Catnb*^+*lox(ex3)*^ transgenic line (Harada et al., 1999) to Acta2-Cre^ERT2^ (SMA-Cre^ERT2^) (Wendling et al., 2009);R26^mTmG^ mice (henceforth referred to as SMACre/Tom and SMACre/β-cat for wild-type and mutant lines, respectively) (Fig.2A). As expected, TAM administration in SMACre/Tom and SMACre/β-cat mice led to GFP expression in the basal cell layer (Fig. 2B, S4A-B). Flow-cytometry analyses 48-72 h after tamoxifen induction confirmed the confinement of SMA-derived GFP+ mammary cells to the basal epithelial compartment (CD24^+^CD29^hi^) in SMACre/Tom control mice (Fig.2C-D). By contrast, a minor percentage of GFP+ cells isolated from SMACre/β-cat mammary glands were also detected in the luminal compartment (CD24^+^CD29^lo^), suggesting perturbation of normal mammary epithelial lineage segregation in response to constitutive β-catenin stabilization within 72 h (Fig.2C-D).

**Fig. 2.**
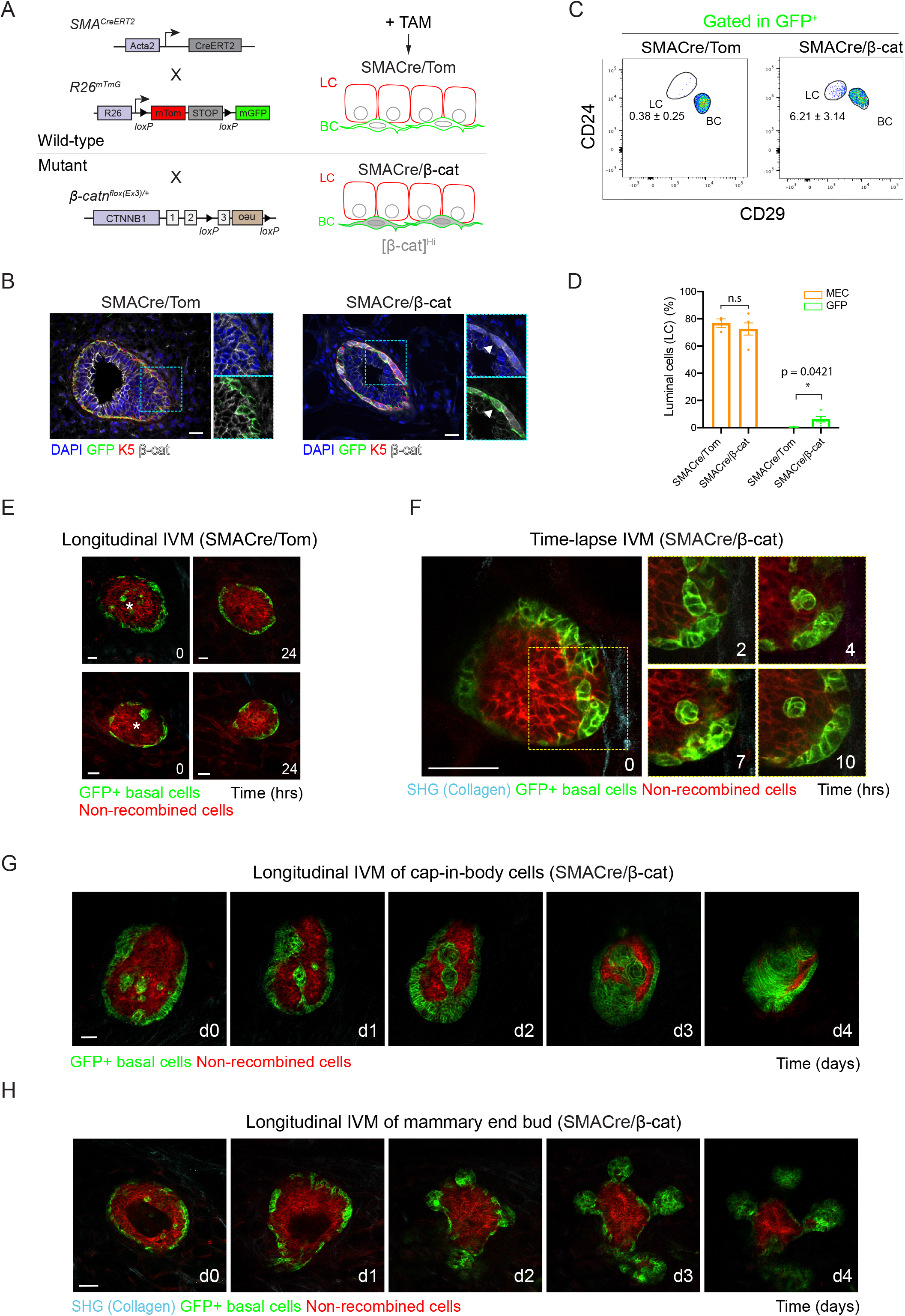
β-catenin activation in mammary basal cells causes aberrant branching morphogenesis and the development of hyperplastic lesions. **(A)** Schematic representation of the smooth muscle actin (SMA)-Cre^ERT2^;R26^mTmG^ (SMACre/Tom) and SMA-Cre^ERT2^;R26^mTmG^;β-catn^flox(Ex3)/+^ (SMACre/β-cat) mouse models used. All cells are labelled with a red membrane tomato fluorescence. Low-dose Tamoxifen (TAM) administration induces membrane GFP labelling of basal cells in both models, and β-catenin accumulation in the SMACre/β-cat model. **(B)** Representative sections of mammary gland tissues showing β-catenin cytoplasmic/nuclear accumulation in K5-expressing basal cells in the mammary epithelium of SMACre/β-cat mice, which coincided with membrane GFP expression (arrowheads). K5 is shown in red, anti-β-catenin staining in white and DAPI labels nuclei in blue. **(C)** Representative FACS dot plots of luminal CD24^+^CD29^lo^ (LC) and basal CD24^+^CD29^hi^ (BC) mammary cells gated within the GFP+ mammary epithelial cell (MEC) population in pubertal SMACre/Tom and SMACre/β-cat mice 48-72 h after tamoxifen administration. While GFP+ MECs are restricted to the basal CD24^+^CD29^hi^ compartment in SMACre/Tom glands, GFP+ MECs from β-cat mutant mice are observed in the CD24^+^CD29^lo^ LC gate. Average values ± SEM are shown. **(D)** Quantification of GFP+ CD24^+^CD29^lo^ cells (green) as compared to the proportion of LCs within MECs (orange) in SMACre/Tom or SMACre/β-cat 48-72 h after tamoxifen administration. Graph shows mean ± SEM (* p = 0.0421, Welch’s t-test, n = 3-5 mice per group). **(E)** IVM images of mammary end bud structures in SMACre/Tom mice showing the elimination of basal GFP+ (green) cap-in-body cells (asterisk) within 24 h. n = 3 mice. **(F)** Acute timelapse IVM of basal GFP+ (green) cap-in-body cell behaviors in a mammary end bud of a pubertal SMACre/β-cat mouse. n = 2 mice. **(G)** IVM images showing the expansion of GFP+ (green) basal cap-in-body cells in terminal end bud structures over time in a pubertal SMACre/β-cat mouse. **(H)** IVM images showing aberrant bud formation over time in the mammary gland of a pubertal SMACre/β-cat mouse. Days, d. (G-H), n = 4 mice. Red: non-recombined membrane tdTomato-expressing mammary epithelial cells in (E-H); cyan: collagen (SHG) in (F-H). d = days in (G-H). Scale bar: 20μm in B; 25μm in E; 50μm in (F-H).

To visualize the impact of β-catenin stabilization on basal cell dynamics, we next performed IVM in SMACre/Tom and SMACre/β-cat pubertal mammary glands 24-72 h after Cre induction. Using this approach, we documented the rapid removal of wild-type GFP+ basal cells (SMACre/Tom) residing in the body cell layer of TEBs (referred to as “cap-in-body” cells) (Fig. 2E, S4C). In contrast, mutant β-catenin GFP+ cap-in-body cells appeared to clonally expand over time (Fig. 2F, Movie S5). To visualize the evolving consequences of constitutive Wnt activation on mammary basal cell dynamics and tissue integrity, we next performed longitudinal IVM in the mammary glands of pubertal SMACre/Tom and SMACre/β-cat mice over time. While wild-type GFP+ basal cells largely retained their typical elongated morphology in mammary ductal structures (Fig.S4D-E), mutant GFP+ cells were cuboidal, and rapidly expanded to generate rosette-like lesions (Fig. 2G, Movie S6). Mutant GFP+ basal cells in apparently normal TEBs and subtending ductal regions were also visualized to swiftly give rise to aberrant bud-like structures, often within 24 h (Fig. 2H). Similar to our observations in N1Cre/β-cat mice (Fig. S3B), GFP fluorescence in the center of lesions gradually diminished over time, suggesting that cells undergo cell death during this process (Fig. 2G-H, Movie S6).

### Constitutive Wnt/β-catenin-induced lesion formation is mediated by hyperproliferation

Acute and longitudinal IVM imaging revealed the dynamic rearrangements and expansion of luminal or basal mammary cells in response to β-catenin stabilization, with similarities observed in the organization of mutant cells during the early stages of lesion development in both models. To investigate the cellular mechanisms underlying the observed phenotypes, we characterized the proliferative capacity of mutant β-catenin cells by 5-ethynyl-2’-deoxyuridine (EdU) incorporation. While rarely detected in the mammary epithelium of wild-type N1Cre/Tom mice (Fig. 3A), the proportion of EdU+ cells was markedly increased in the mammary glands of N1Cre/β-cat mice, even within morphologically normal-looking mammary ducts (Fig. 3A-B). Surprisingly, increased EdU uptake was observed in both luminal and basal compartments prior to lesion development, suggesting a non-cell autonomous effect (Fig. 3B). EdU incorporation correlated with β-catenin accumulation in morphologically normal-looking ducts, nascent luminal lesions, and more advanced aberrant structures (Fig. 3C), further indicating that increased proliferation contributed to the cellular dynamics observed by intravital imaging.

**Fig. 3.**
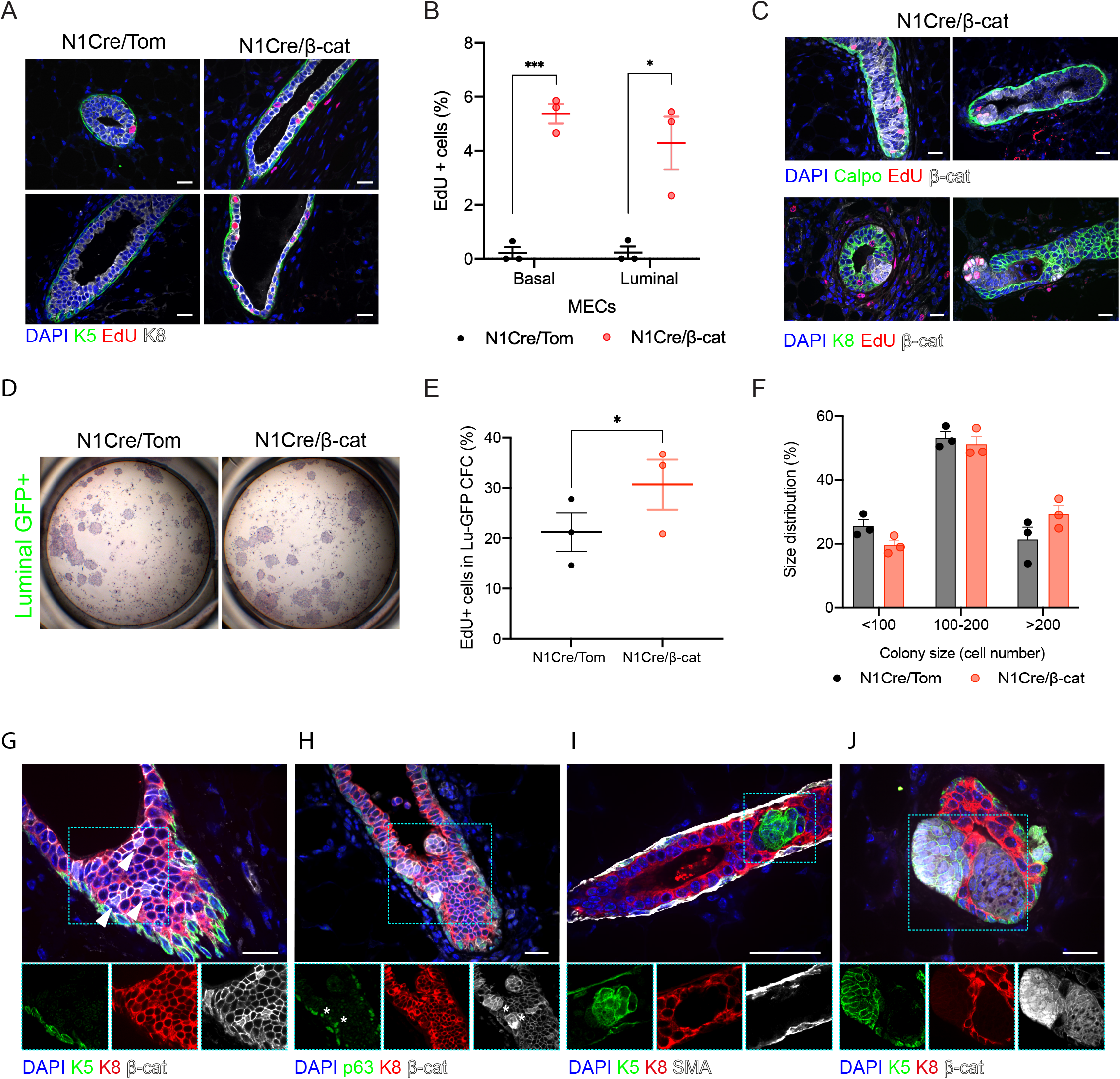
β-catenin stabilization in mammary luminal cells leads to increased proliferation and the development of lesions with aberrant lineage marker expression. **(A)** Representative sections of mammary gland tissues from N1Cre/Tom and N1Cre/β-cat mice showing EdU+ (red) cells in the luminal (K8, white) and basal (K5, green) epithelial compartment. (**B)** Quantification of the percentage of EdU+ (GFP+ or GFP-) cells in the basal and luminal mammary epithelial cell (MEC) compartment in phenotypically normal ductal structures in wild-type and mutant β-catenin models. Graph shows mean ± SEM (* p < 0.05, *** p = <0.001 Welch’s t-test; n = 3 animals per group, 3 weeks after TAM induction). **(C)** Edu+ cells (red) in mutant N1Cre/β-cat mammary sections coincided with β-catenin accumulation (white) in nascent and more established lesions 6-7 days after TAM induction. **(D)** No differences were observed in the colony forming capacities of GFP+ luminal cells isolated by flow cytometry from the mammary glands of N1Cre/Tom and N1Cre/β-cat mice 3 weeks after TAM induction. Representative images of hematoxylin and eosin-stained colonies after 7 days in culture. **(E)** Colonies arising from GFP+ luminal cells isolated from N1Cre/β-cat mice possessed a higher percentage of proliferative (EdU+) cells. Graph shows mean ± SEM (* p < 0.05, paired t-test; n=3 independent sorting experiments). **(F)** Size distribution of colonies generated from isolated wildtype and mutant GFP+ luminal cells. Graph shows mean ± SEM (p < 0.05, Pearson’s chi-square test; n=3 independent sorting experiments performed 3 weeks after induction). **(G-J)** Representative sections of mammary ducts immunostained for β-catenin (white) in N1Cre/β-cat mice showing changes in lineage marker expression with hyperplastic lesion development. β-catenin accumulation (marked by white arrows) in luminal K8 expressing cells (red) in a phenotypically normal duct 6 days after TAM induction **(G)**. The asterisks in **(H)** show aberrant expression of basal marker p63 (green) in early mutant β-catenin luminal K8+ lesions 6-7 days after TAM induction. **(I-J)** Larger mutant β-catenin derived lesions are devoid of K8 (red) expression and express K5 (green), a mammary basal marker in mammary tissues harvested 3 weeks after TAM induction. Scale bars: 20 μm.

To investigate if β-catenin activation affected the clonogenic capacity of Notch1-expressing luminal progenitors, GFP+ luminal cells from N1Cre/Tom and N1Cre/β-cat mice were isolated by flow cytometry and seeded on a feeder layer of irradiated 3T3 fibroblasts (Sleeman et al., 2007). No differences were observed between the colony-forming potential of wild-type and mutant β-catenin mammary luminal cells, suggesting that progenitor cell frequencies were comparable between the two models (Fig. 3D, Fig.S5A). Mutant colonies contained a higher proportion of proliferating (EdU+) cells, however, generating larger colonies and corroborating our *in vivo* observations (Fig. 3E-F).

We next sought to phenotypically characterize the lesions arising in response to luminal targeting of β-catenin in N1Cre/β-cat mice. While restricted to basolateral membranes in wild-type mammary cells (Fig.S5B), β-catenin accumulation was clearly visible in the cytoplasm of K8-expressing luminal cells in the mammary glands of N1Cre/β-cat mice shortly after Cre induction (Fig.3G). β-catenin stabilization was concentrated in nascent luminal lesions that retained luminal marker expression, although rare p63-expressing luminal (K8+) cells were also observed (Fig. 3H, S5C). Over 3-4 weeks, certain Notch1-derived lesions appeared to lose K8 expression and gain expression of the basal cytokeratin K5 (Fig. 3I-J, S5C), suggesting the acquisition of basal characteristics in response to Wnt activation. Of interest, these lesions invariably lacked the expression of the basal cell differentiation marker SMA (Fig. 3I, S5C). Moreover, clusters of basally-located, K5-expressing cells with stronger nuclear β-catenin staining were frequently observed next to inner cell clusters that exhibited weaker staining (Fig. 3J). Lesions eventually developed into large rosette-like structures that encircled islands of keratin debris and cells with aberrantly shaped or absent nuclei (“ghost cells”) (Fig.S5D), consistent with previous observations of constitutive Wnt/β-catenin activation (Jardé et al., 2016; Miyoshi et al., 2002b, 2002a). Terminal transferase-mediated dUTP nick end labelling (TUNEL) assays confirmed that cells within rosette-like lesions undergo cell death during this process (Fig. S5E).

Analogous EdU incorporation studies in SMACre/Tom and SMACre/β-cat mice also revealed a marked increase in the proportion of EdU+ basal cells prior to lesion development (Fig. 4A). Notably, similar to N1Cre/β-cat mice, increased proliferation was also detectable in luminal cells in morphologically normal ductal structures, indicative of paracrine signaling (Fig. 4B). The high rate of recombination induced by *SMA-Cre^ERT2^* enabled us to quantify the proportion of proliferating GFP+ cells in wild-type and mutant models, revealing a 10-fold increase in the mutant ductal epithelium, rising to nearly 30-fold in visible lesions (Fig. 4C). EdU incorporation was particular evident in cells with β-catenin accumulation in mutant-derived lesions (Fig. 4D). In line with our results with luminal targeted cells, β-catenin activation had no significant impact on the *in vitro* colony-forming efficiency of basal GFP+ cells (Fig.4E, Fig.S6A), however mutant colonies were more proliferative and significantly larger than control colonies (Fig.4E-G).

**Fig. 4.**
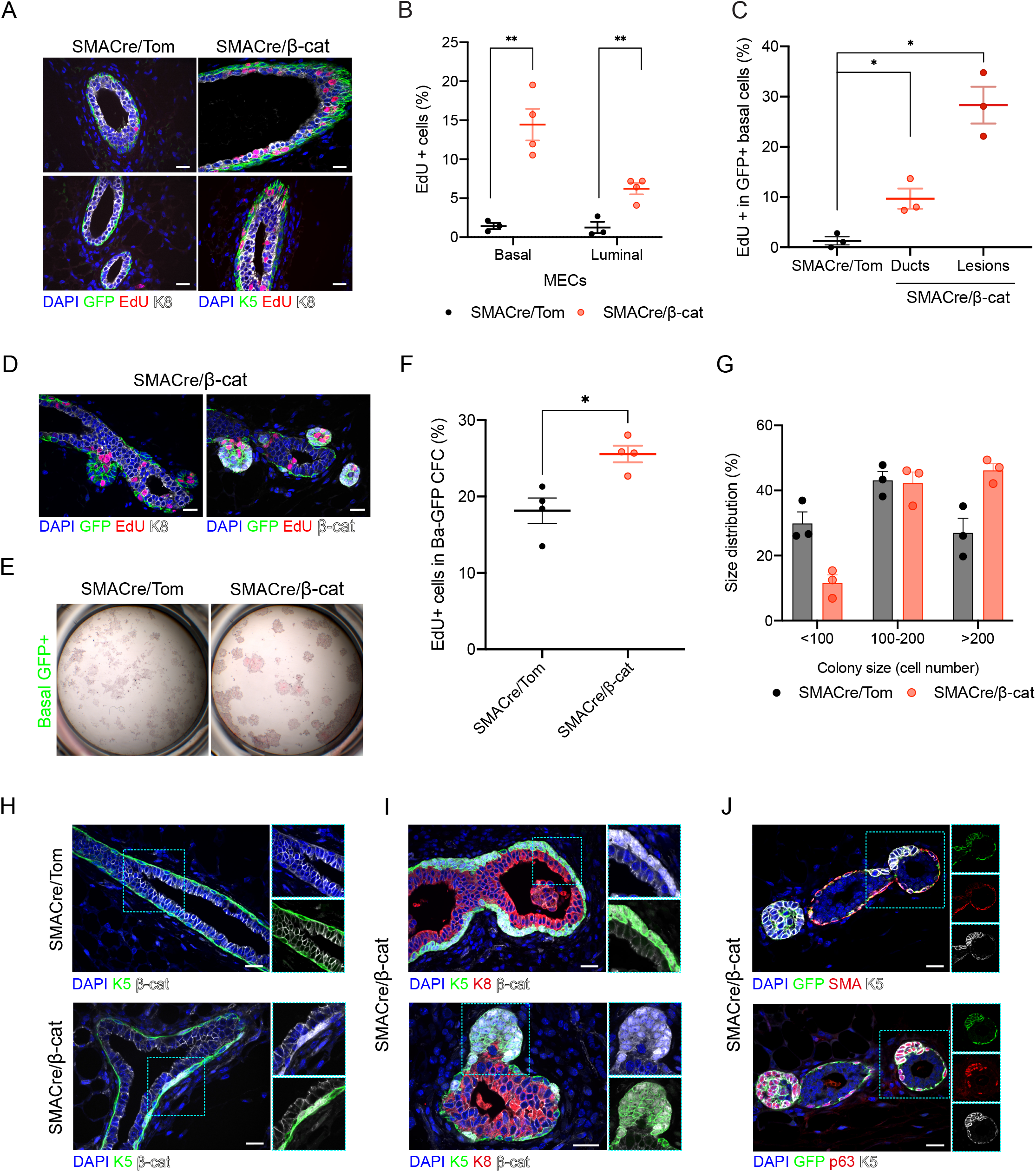
β-catenin stabilization in mammary basal cells leads to increased proliferation and the development of lesions with aberrant lineage marker expression. **(A)** Representative sections of mammary glands from SMACre/Tom and SMACre/β-cat mice showing EdU+ (red) cells in the luminal (K8, white) and basal (K5 or GFP, green) compartments. (**B)** Quantification of the percentage of EdU+ cells in basal and luminal mammary epithelial cells (MECs) in phenotypically normal ductal structures in SMACre/Tom and SMACre/β-cat mice. Graph shows mean ± SEM (** p < 0.01 Welch’s t-test; n=3 animals per group). **(C)** Quantification of proliferative GFP+ basal cells in SMACre/Tom mammary sections, normal ducts and aberrant regions in SMACre/β-cat mammary tissue. Graph shows mean ± SEM (* p < 0.05, Welch’s t-test; n=3). **(D)** Edu+ cells (red) in mutant SMACre/β-cat mammary tissue sections coincided with GFP expression and β-catenin accumulation (white) in mammary lesions. **(E)** No statistically significant differences were observed in the colony forming capacities of GFP+ basal cells isolated by flow cytometry from the mammary glands of SMACre/Tom and SMACre/β-cat mice. Representative images of hematoxylin and eosin-stained colonies after 7 days in culture. **(F)** Colonies arising from GFP+ basal cells isolated from SMACre/β-cat mice present a higher percentage of proliferative (EdU+) cells. Graph shows mean ± SEM (* p < 0.03, paired t-test; n=4 independent sorting experiments). **(G)** Size distribution of colonies generated from isolated wildtype and mutant GFP+ luminal cells. Graph shows mean ± SEM. p < 0.0001, Pearson’s chi-square test; n=3 independent sorting experiments. **(H)** Representative sections of mammary ducts showing β-catenin cytoplasmic/nuclear accumulation in keratin(K)5-expressing basal cells in the mammary epithelium of SMACre/β-cat mice. **(I)** Representative images of mammary sections showing β-catenin-induced changes to basal cell morphology and the development of aberrant bud-like lesions in SMACre/β-cat mice. **(J)** Aberrant basal-derived lesions express K5 and p63 but lack SMA expression. All images were acquired in tissues harvested 3-7 days after TAM induction. Scale bars: 20 μm.

While no cytoplasmic/nuclear β-catenin could be observed in basal cells of SMACre/Tom mice, SMACre/β-cat basal cells exhibited strong nuclear β-catenin staining and a more cuboidal morphology compared to wild-type basal cells (Fig.4H-I). Although lesions retained the expression of the basal lineage markers K5 and p63, they invariably lacked SMA expression, a marker for myoepithelial differentiation (Fig.4I-J). As observed by IVM (Fig. 2), mutant growths rapidly evolved into large rosette-like, keratinized structures containing ghost cells lacking mammary epithelial marker expression (Fig. S6B), eventually undergoing cell death (Fig. S6C).

### Constitutive Wnt/β-catenin signaling induces squamous transdifferentiation of mammary epithelial cells

Our longitudinal IVM studies and histological analysis of early mutant β-catenin-induced lesion development consistently revealed the formation of stereotypical bud-like clusters of epithelial cells (Fig.S7A). These frequently displayed divergent upward displacement of epithelial nuclei towards newly forming growths, with epithelial cells organizing themselves into a compact arrangement (Fig.S7A). This epithelial clustering into ring-like structures is reminiscent of early stage embryonic Hair Follicle (HF) formation (Devenport and Fuchs, 2008), and that observed in adult mouse skin in response to β-catenin activation in hair follicle stem cells, which drives new axes of hair follicle growth (Deschene et al., 2014). Moreover, long-term, non-lineage-specific targeting of stabilized β-catenin in mammary tissues using MMTV-Cre and WAP-Cre models was previously shown to induce squamous metaplasia, a process characterized by the expression of epidermal markers (Miyoshi et al., 2002b, 2002a). Considering these similarities, we next investigated whether luminal or basal targeting of mutant β-catenin resulted in epidermal transdifferentiation. Immunostaining for the HF marker K6 - normally only expressed in TEBs during puberty (Grimm et al., 2006; Smith et al., 1990) - revealed its high expression in lesions in both N1Cre/β-cat and SMACre/β-cat models (Fig. S7B-D). At more advanced stages, these structures exhibited high hair keratin (HK) expression (Fig. S7C-D), although no detectable expression of the suprabasal skin marker K10 was observed (Fig. S7C-D). Collectively, these findings suggest that, regardless of the mammary lineage targeted, β-catenin stabilization drives the acquisition of a HF epidermal differentiation program in mammary epithelial cells, eventually resulting in squamous metaplasia (Fig.S8).

## Discussion

Dysregulated Wnt/β-catenin activity is a hallmark of several types of cancer. Links between Wnt signaling and mammary tumorigenesis are well-established, yet its role in the initiation of different breast cancer subtypes remains poorly understood (van Schie and van Amerongen, 2020). The early molecular and cellular mechanisms underlying Wnt-driven mammary tumorigenesis are particularly unclear, largely due to a reliance on analyzing advanced tumors in previous studies. This is further hampered by discrepancies observed between available transgenic models of Wnt pathway activation, and the different mammary cell lineages targeted (Imbert et al., 2001; Miyoshi and Hennighausen, 2003; Miyoshi et al., 2002b, 2002a; Moumen et al., 2013; Teissedre et al., 2009; Teuliere et al., 2005). Moreover, until now, the impact of targeting a stabilized form of β-catenin refractory to degradation from its endogenous promoter (using *Catnb*^+*lox(ex3)*^ transgenic mice (Harada et al., 1999)) to the mammary basal epithelial compartment, and how this compares to the phenotype observed when targeted to hormone responsive luminal cells remained unexplored. Herein, we sought to assess the phenotypes elicited by the same oncogenic β-catenin mutant on the *in situ* behaviors of distinct mammary epithelial cell types, by coupling longitudinal IVM with lineage specific genetic activation of the Wnt/β-catenin pathway.

Real-time, *in vivo* imaging by IVM provides an essential window into the dynamic cellular events contributing to tissue development and pathology. This powerful approach has provided important insights into the growth, progression, metastasis and therapeutic responses of a plethora of cancer types, including breast cancer (Condeelis and Weissleder, 2010; Ellenbroek and Van Rheenen, 2014; Lloyd-Lewis, 2020). Yet, the application of IVM to study the earliest stages of neoplastic development in the mammary gland remains largely absent, with most previous studies focused on imaging established tumors (Lloyd-Lewis, 2020). Here, we were able to capture the earliest cellular events underlying the impact of mutagenic Wnt/β-catenin signaling on the dynamics of distinct subpopulations of mammary epithelial cells *in vivo* and in real time, and its effects on the organization and differentiation of the mammary epithelium on a tissue scale. Interestingly, IVM revealed that constitutive stabilization of β-catenin from its endogenous promoter in either luminal or basal mammary lineages caused epithelial cells to cluster into ring-like structures, forming compact, bud-like growths that rapidly expanded over time. This was particularly evident when targeted to the basal mammary epithelial layer, where precocious budding could be readily visualized by day-to-day IVM. This behavior showed remarkable similarities to that observed during ectopic HF formation in response to β-catenin activation in HF stem cells (Deschene et al., 2014). Indeed, our data revealed that these early, β-catenin-induced changes to epithelial cell organization and behavior reflected the progressive transdifferentiation of mammary cells to a hair follicle/epidermal-like fate, marked by the acquisition of K6 and HK marker expression (Fig.S8). Based on our findings, we believe that constitutive Wnt activity in mammary luminal cells first drives their conversion to basal-like cells, by repressing K8 expression and acquiring basal traits such as K5 and p63 expression, and that they subsequently enter the same program of squamous transdifferentiation as mutant β-catenin mammary basal cells. This may underlie the observed differences in the timing and rate of keratinized lesion development in response to lineage-specific mutant β-catenin activity, with the appearance of rosette-like structures in the mammary epithelium of N1Cre/β-cat mice taking considerably longer as compared to SMACre/β-cat mice. However, intrinsic differences in recombination efficiency between the two inducible Cre lines used in this study could also account for the different latencies in lesion development.

Finally, our data, consistent with published work (Miyoshi et al., 2002b, 2002a), revealed that the aberrant mammary differentiation program induced by β-catenin activation – evident when targeted to either of the mammary epithelial lineages – leads to the formation of squamous metaplasia composed of keratinized “rosette-like’” structures and the appearance of ghost cells (characteristic of epidermal lesions). This is in contrast to alternative transgenic lines whereby N-terminally truncated β-catenin is ectopically expressed downstream of mammary lineage gene promoters, which give rise to adenocarcinomas (Imbert et al., 2001; Michaelson and Leder, 2001; Teissedre et al., 2009; Teuliere et al., 2005). Discrepancies between the models may be explained by potential inherent differences between transgenic lines, in addition to differences in the potency of β-catenin activation. Indeed, levels of accumulated β-catenin driven by its endogenous promoter in the mammary glands of *Catnb*^+*lox(ex3)*^ transgenic mice (Harada et al., 1999) may be insufficient to drive adenocarcinoma development, compared to N-terminally truncated β-catenin transgenes driven by strong promoters such as a MMTV-LTR (Imbert et al., 2001; Michaelson and Leder, 2001; Miyoshi and Hennighausen, 2003).

In summary, by high-resolution longitudinal IVM, we visualized the dynamic cellular changes induced by lineage-specific mutant β-catenin accumulation *in situ* during the earliest stages of hyperplastic lesion development. We show that, regardless of the targeted cell of origin, aberrant β-catenin accumulation induces a hair-follicle/epidermal differentiation program in mammary epithelial cells, leading to the formation of squamous metaplasia. Importantly, our findings provide further evidence that the mammary epithelium is inherently predisposed towards acquiring an epidermal-like fate upon constitutive Wnt/β-catenin activation. As similar phenotypes are observed in other glandular tissues, including the prostate (Bierie et al., 2003), aberrant Wnt/β-catenin pathway activity is likely capable of over-riding existing cell intrinsic differentiation program in several epithelial cell types to drive pathological tissue development.

## Materials and Methods

### Ethics Statement

All studies and procedures involving animals were in strict accordance with the recommendations of the European Community (2010/63/UE) for the Protection of Vertebrate Animals used for Experimental and other Scientific Purposes. Approval was provided by the ethics committee of the Institut Curie CEEA-IC #118 and the French Ministry of Research (reference #04240.03). We comply with internationally established principles of replacement, reduction, and refinement in accordance with the Guide for the Care and Use of Laboratory Animals (NRC 2011). Husbandry, supply of animals, as well as maintenance and care in the Animal Facility of Institut Curie (facility license #C75–05–18) before and during experiments fully satisfied the animal’s needs and welfare. All animals were housed in individually ventilated cages under a 12:12 h light-dark cycle, with water and food available ad libitum. All mice were sacrificed by cervical dislocation.

### Mouse models

All mouse lines used have been previously described and were of mixed genetic background. N1Cre^ERT2^ (Fre et al., 2011) and SMACre^ERT2^ (Wendling et al., 2009) were crossed to the double fluorescent reporter Rosa26^mT/mG^ (Muzumdar et al., 2007) and inducible mutant β-catenin (Harada et al., 1999) (kindly provided by Lionel Larue, Institut Curie) transgenic lines. We exclusively analysed female mice and no randomization methods were performed. Reporter expression and β-catenin stabilisation was induced in N1Cre^ERT2^ or SMACre^ERT2^ females by a single intraperitoneal injection of tamoxifen free base (Euromedex) prepared in sunflower oil containing 10% ethanol (0.1 mg per g of mouse body weight), unless indicated otherwise in figure legends. Using this dose, mammary gland development appeared to progress unabated, as previously reported (Rios et al., 2014). In experiments exceeding 8-10 days, tamoxifen doses were reduced 10-fold (0.01mg per g of mouse body mice) to avoid potential systemic toxicity following long-term β-catenin stabilisation. For EdU (5-ethynyl-2-deoxyuridine) labelling experiments, mice were injected intraperitoneally with 20 mg per kg body weight of EdU 2 h before harvesting mammary gland tissues.

### Intravital imaging

Intravital imaging through a titanium/glass mammary imaging window were based on previously published protocols (Messal et al., 2021). Intravital imaging through our custom-made PDMS imaging windows was performed as previously described (Jacquemin et al., 2021). Briefly, mammary imaging windows were surgically implanted over the fourth mammary glands of 5-6 week mice at the indicated times after tamoxifen administration. Mice were anaesthetized using isoflurane (1.5% isoflurane/medical air mixture) and placed in a facemask with a custom designed holder to stabilize windows during imaging acquisition. Imaging was performed on an upright Nikon A1R multiphoton microscope equipped with a Spectra-Physics Insight Deepsee laser, conventional and resonant scanners, GaAsP non-descanned detectors using 16x NA 0.8 or 25x NA 1.1 PlanApo LambdaS water objectives. An excitation wavelength of 960 nm was used for GFP and TdTomato, in addition to second harmonic generation (SHG) imaging of collagen. Mammary epithelial structures imaged in timelaspe were acquired every 30 min using a *Z*-step size of 2 μm. For long-term, longitudinal imaging z-stacks (with z-step size of 2 μm) of epithelial structures were taken either 1-2x daily for up to 8 days, or 2-3x weekly for up to 3 weeks (as indicated in figure legends).

For high-resolution reconstruction of time-lapse and longitudinal acquisitions, regions of interest encompassing discrete Z-stack sizes were selected and registered using the StackReg Rigid Body plugin in FIJI (ImageJ v1.53) (Long et al., 2012). Images were processed using a Gaussian blur filter (0.5-1 pxl radius) in FIJI (ImageJ v1.53).

### Optical tissue clearing and whole-mount immunostaining

Mammary gland pairs 2 or 3 were dissected, spread on TetraPak card and optimally fixed for 6-9 h in 10% Neutral Buffered Formalin (NBF) at room temperature. Mammary glands were cut into large pieces (~15×15×2 mm) for immunostaining and tissue clearing, as previously described in detail (Lloyd-Lewis et al., 2016). Optical tissue clearing was performed using a modified CUBIC (Reagent 1A) protocol (Susaki and Ueda, 2016). Briefly, tissues were immersed in CUBIC Reagent 1A (urea (10% w/w), Quadrol^®^ (5% w/w), triton X-100 (10% w/w), NaCl (25 mM) in distilled water] for 2-3 days at 37°C, washed in PBS and blocked overnight at 4°C in PBS containing normal goat serum (10%) and triton X-100 (0.5%). Tissue was incubated in primary antibodies diluted in blocking buffer at 4°C for 4 days with gentle agitation. Tissue was washed in PBS (3 x 1 h) and incubated with secondary Alexa-fluor conjugated antibodies at 4°C for 2 days with gentle agitation before further washing in PBS (3 x 1h) and incubation with DAPI (10 μM) for 2-3 h at room temperature. Tissues were imaged in CUBIC Reagent 2.

### Immunohistochemistry

IHC was performed according to a previously published protocol (Sargeant et al., 2014). Briefly, formalin-fixed paraffin embedded 5-7 μm mammary tissue sections were deparaffinized in xylene and rehydrated in a reducing ethanol series. Tissue was permeabilized in phosphate buffered saline (PBS) containing triton X-100 (0.5%). Heat-induced epitope retrieval was performed in sodium citrate (0.01 M, pH 6) for 11 min at 110°C. Slides were blocked in PBS containing fetal bovine serum (5%) BSA (2%) and triton x-100 (0.25%) for 1 h. Primary antibodies diluted in blocking buffer were incubated overnight at 4°C in a humidified chamber. Secondary antibodies diluted in PBS were incubated for 1 h at room temperature. EdU detection was performed using the Click-iT EdU Alexa Fluor 647 Imaging Kit (Molecular Probes), according to the manufacturer’s instructions. Nuclei were stained with DAPI dilactate (625 ng/mL) for 10-15 min at room temperature or, in the case of Edu detection, with Hoechst33342 10 μg/ml for 30 min at room temperature. Slides were mounted using Aqua-Polymount.

### Fluorescence confocal microscopy of fixed tissues

#### 3D imaging

CUBIC-cleared tissues were imaged in CUBIC Reagent 2 in 35 mm glass-bottom Fluoro-dishes. Images were acquired using an LSM780 or LSM880 inverted laser scanning confocal microscope (Carl Zeiss) equipped with 10×/0.3 PL NEOFLUAR or 25×/0.8 LD LCI PLAN-APO objective lenses. For standard 4-colour imaging, laser power and gain were adjusted manually to give optimal fluorescence for each fluorophore with minimal photobleaching. Imaging depths were recorded from the top of the epithelial structure being imaged (typically ~350 μm through the native fat pad). Image reconstructions were generated in FIJI (ImageJ v1.53) using the Bio-Formats plugin (National Institutes of Health) (Linkert et al., 2010; Schindelin et al., 2012). Denoising of 3D image stacks was performed in MATLAB (R2014a, The Mathoworks Inc.) (Boulanger et al., 2010).

#### 2D imaging

Images of stained sections were acquired using an upright spinning disk (CSU-X1 scan-head from Yokogawa) confocal microscope (Carl Zeiss, Roper Scientific France), equipped with a CoolSnap HQ2 charge coupled device (CCD) camera (Photometrics) and PLAN APO ×63/1.4 NA and PLAN APO 40x/1.3NA objective lenses. Images were captured using Metamorph and processed in FIJI (ImageJ v1.53).

### Mammary gland dissociation and Flow Cytometry

Single cell dissociation was performed through enzymatic digestion with 600 U ml^−1^ collagenase (Sigma) and 200 U ml^−1^ hyaluronidase (Sigma) for 90 min at 37 °C. Cells were further dissociated in TrypLE (Gibco) for 3 min, in 5 mg ml^−1^ dispase (Roche) and 0.1 mg ml^−1^ DNase I (Sigma) for 5 min, and then in 0.63% NH_4_Cl and filtered through a 40 μm cell strainer to obtain a single cell preparation for FACS. Cell labelling and flow cytometry were performed as described previously (Koren et al., 2015) using LSRII or FACS ARIA flow cytometers (BD). Dead cells (DAPI^+^), and CD45^+^/CD31^+^/Ter119^+^ (Lin^+^) non-epithelial cells were excluded. The following antibodies were all purchased from Biolegend and were used at a 1:100 final concentration: PE/Cy7 anti-mouse CD24, PE/Cy7 anti-mouse Epcam, AlexaFluor700 anti-mouse/rat CD29, APC/Cy7 anti-mouse CD49f, lineage markers: APC anti-mouse CD31, APC anti-mouse Ter119, APC anti-mouse CD45, isotype controls: PE rat IgM, PerCP/Cy5.5 rat IgGa, PE/Cy7 rat IgG2a, APC/Cy7 rat IgG2a, APC rat IgG2b. The purity of sorted populations was approximately 95%. The results were analysed using FlowJo software (v10, BD).

### Colony forming assays

Sorted cells well cultured on irradiated 3T3 cell feeders in 24-well plates for 7 days. Luminal cells were plated at a density of 400 cells per well and cultured in DMEM/F12 medium containing fetal bovine serum (10%), 5 μg/ml insulin (Sigma-Aldrich), 10 ng/ml EGF (Invitrogen, Life Technologies) and 100 ng/ml cholera toxin (ICN Biochemicals). Basal cells were plated at a density of 1500 cells per well and cultured in DMEM/F12 medium containing fetal bovine serum (1%), 1/50 diluted B27 supplement (ThermoFisher), 5 μg/ml insulin (Sigma-Aldrich), 10 ng/ml EGF (Invitrogen, Life Technologies). Colonies were fixed in 4% PFA and stained with hematoxylin/eosin and pictures were acquired using a Leica MZ8 binocular. To evaluate colony number and size, ImageJ software was used. For proliferation analysis, colonies were incubated with EdU at 2 μg/ml for 1 h prior to PFA fixation. Edu detection was performed with Click-iT EdU Alexa Fluor 647 Imaging Kit (Molecular Probes), according to the manufacturer’s instructions.

### Statistics and Reproducibility

Experiments were performed in biological replicates as stated in figure legends. For each experiment, we have used at least n=3 animals, and experiments with at least n=3 replicates were used to calculate the statistical value of each analysis. Data processing and statistical analysis were performed in Prism (v9, GraphPad). All graphs show mean ± SEM. Statistical analysis was performed with two-tailed unpaired Welsh’s t-tests, unless otherwise stated in figure legends.

---

**Table S1.** Reagents and materials.

| Reagents | Supplier | Reference |
| --- | --- | --- |
| 10% NBF | Sigma | HT501128 |
| Triton X-100 | Euromedex | 2000-C |
| 2,2',2''-nitrilotriethanol | Sigma | 90279 |
| Quadrol | Sigma | 122262 |
| Urea | Sigma | U5378 |
| Sucrose | Sigma | S0389 |
| Sunflower Oil | Sigma | S5007 |
| Tamoxifen | MP Biomedicals | 156738 |
| Materials | Supplier | Reference |
| 35mmglass bottom dishes | Fluorodish | 81158 |

**Table S2.** Antibodies.

| Antibody | Dilution (2D) | Dilution (3D) | Supplier | Reference |
| --- | --- | --- | --- | --- |
| rabbit anti-SMA | 1:200 | 1:300 | Abcam | ab5694 |
| mouse anti-SMA | 1:200 | - | Sigma-Aldrich | A2547 |
| rat anti-K8 | 1:200 | 1:50 | Developmental Studies Hybridoma Bank | TROMA-I |
| mouse anti-Beta-catenin | 1:50 | - | BD | 610154 |
| rabbit anti-Keratin 6 | 1:200 | - | Covance | PRB-169P |
| mouse anti-Keratin 10 | 1:100 | - | Dako | M7002 |
| rabbit anti-keratin 5 | 1:300 | - | Covance | PRB-160P |
| chicken anti-GFP | 1:300 | - | Novus Biologicals | NB 100-1614 |
| goat anti-rabbit AlexaFluor (AF) 647 | 1:500 | 1:500 | Thermo Fisher Scientific | A21245 |
| goat anti-rat AlexaFluor (AF) Cy5 | 1:500 | - | Thermo Fisher Scientific | A10525 |
| goat anti-rabbit AlexaFluor (AF) Cy3 | 1:500 | - | Thermo Fisher Scientific | A10520 |
| goat anti-rabbit<br>AlexaFluor (AF)<br>488 | 1:500 | - | Thermo Fisher<br>Scientific | A21206 |
| goat anti-mouse<br>AlexaFluor (AF)<br>Cy5 | 1:500 | - | Thermo Fisher<br>Scientific | A10524 |

## Acknowledgements

The authors thank P. Chambon and D. Metzger for providing the SMACre^ERT2^ (Acta2-Cre^ERT2^) mice, S. Tajbakhsh for the mTmG reporter line, and L. Larue for sharing the β-catenin floxed line. The authors acknowledge the Flow Cytometry and Cell Sorting Platform at Institute Curie for their expertise; the In Vivo Experimental Facility for help in the maintenance and care of our mouse colony; and the Experimental Pathology facility at Curie Hospital for paraffin sample preparation. The authors especially thank Lucie Sengmanivong and Marie Irondelle for intravital imaging support. The PICT-IBiSA imaging platform was funded by ANR-10-INBS-04 (France-BioImaging), ANR-11 BSV2 012 01, ERC ZEBRATECTUM no. 311159, ARC SFI20121205686 and the Schlumberger Foundation. This work was supported by Paris Sciences et Lettres (PSL* Research University), the French National Research Agency (ANR) grant (no. ANR-15-CE13-0013-01), the Schlumberger Foundation and by Labex DEEP ANR-Number 11-LBX-0044. B.L-L is also funded by a Vice-Chancellor’s Research fellowship from the University of Bristol, and support from The Academy of Medical Sciences. The funders had no role in study design, data collection and analysis, decision to publish, or preparation of the manuscript.

## Author Contributions

B.L-L, M.M.F and S.F. conceived and designed the experiments. B.L-L, M.P, F.G, G.J, M.M.F performed all experiments and analysis. B.L-L and S.F. wrote the manuscript. All authors reviewed and approved the manuscript.

## Data availability

Data supporting the findings of this study are available within the article. Requests for further information should be directed to the corresponding author.

