## Supplementary Figures for "*In vivo* imaging of mammary epithelial cell dynamics in response to lineage-biased Wnt/β-catenin activation"

**This PDF file includes:**

Figs. S1 to S8
Captions for Supplemental Movies S1 to S6

**Other Supplemental Materials for this manuscript include the following:**

Movies S1 to S6

Supplementary Figure 1

A

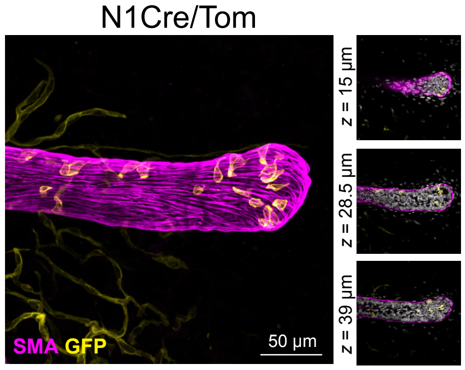


B

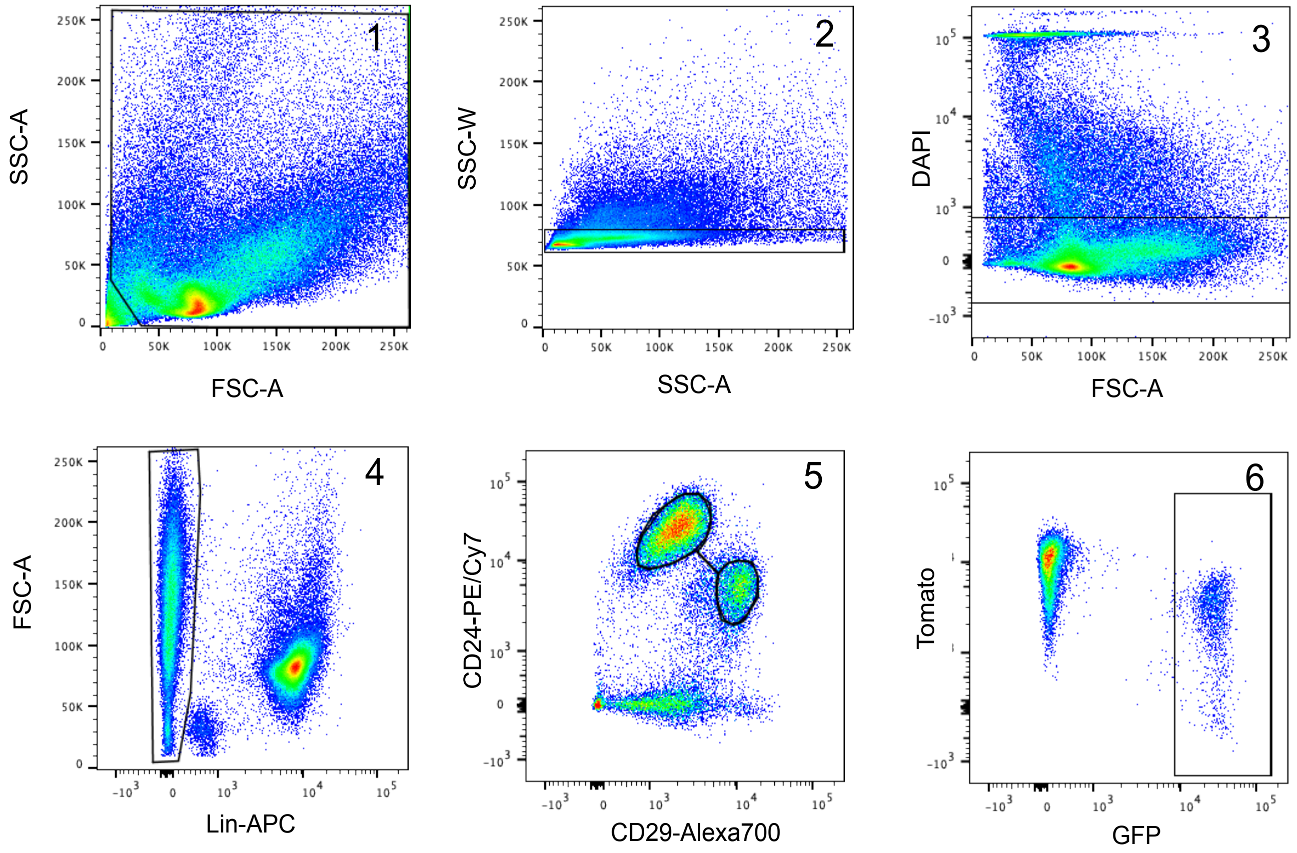

**Fig. S1. Wholemount imaging and flow cytometry analysis of mammary tissues. Related to Fig. 1.**

**(A)** Maximum intensity z-projection and thin optical slices (depth in *z* relative to the first image in the image sequence) of cleared mammary tissue from N1Cre/Tom mice immunostained with smooth muscle actin (SMA in pink). GFP+ cells (shown in yellow) are restricted to the luminal compartment < 1 week after low-dose tamoxifen administration. Scale bar: 50μm. **(B)** Representative dot plots of the applied FACS gating strategy. (1) FCS/SSC gating allows cell debris to be discarded, (2) SSC-A/SSC-W selects single cells, (3) DAPI exclusion selects live cells, (4) Lin exclusion (CD45/CD31/Ter119 cell surface marker proteins) eliminates hematopoietic cells, (5) CD29/CD24 cell surface markers are used to identify Mammary Epithelial Cells (MEC) and (6) GFP/Tomato selects fluorescent cells.

### Supplementary Figure 2

A

Longitudinal IVM of a TEB during pubertal mammary development (N1Cre/Tom)

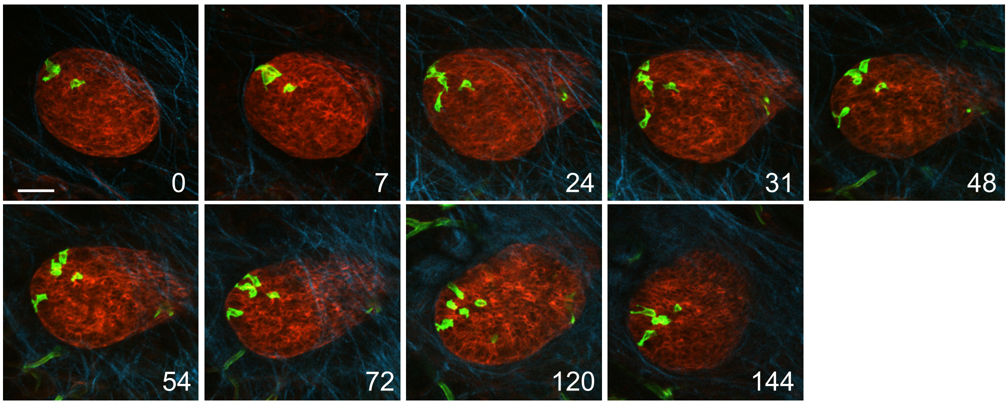


SHG (Collagen) GFP+ luminal cells Non-recombined cells Time (hrs)

B

Longitudinal IVM of a duct during pubertal mammary development (N1Cre/Tom)

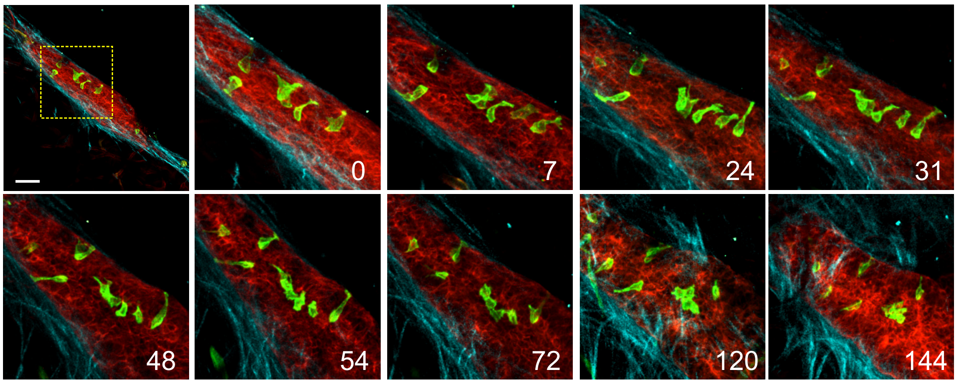


SHG (Collagen) GFP+ luminal cells Non-recombined cells Time (hrs)

C

Longitudinal IVM of ductal lumen formation

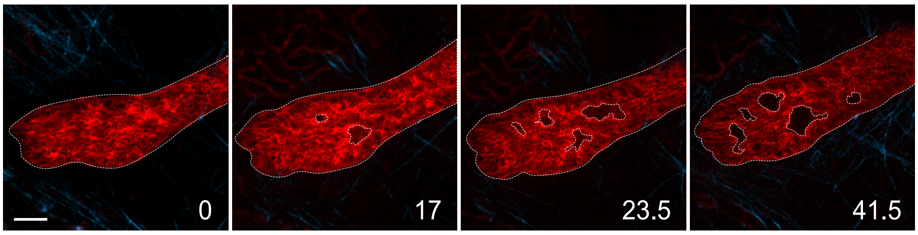


SHG (Collagen) MECs Time (hrs)

D

Longitudinal IVM of end bud lumen formation

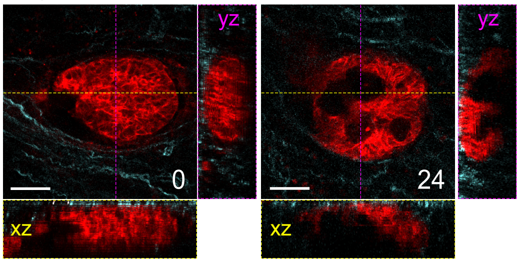


SHG (Collagen) MECs Time (hrs)

**Fig. S2. Longitudinal IVM of wildtype luminal cell behaviors during pubertal ductal development. Related to Fig.1.**

**(A-B)** IVM images of luminal GFP+ (green) mammary cells in a mammary end bud structure (A) or in a subtending duct (B) of a pubertal N1Cre/Tom mouse over time. Close up images included in Figure 1F, related to Supplemental Movie 1. **(C)** IVM images of a mammary ductal structure showing the process of lumen formation. **(D)** Longitudinal intravital images of a mammary TEB showing rapid lumen formation. Orthogonal views show *XZ* (*yellow line and box*) and *YZ* (*purple line and box*) planes. Red: non-recombined membrane tdTomato-expressing mammary epithelial cells; cyan: collagen (SHG). All scale bars: 50μm.

Supplementary Figure 3

A

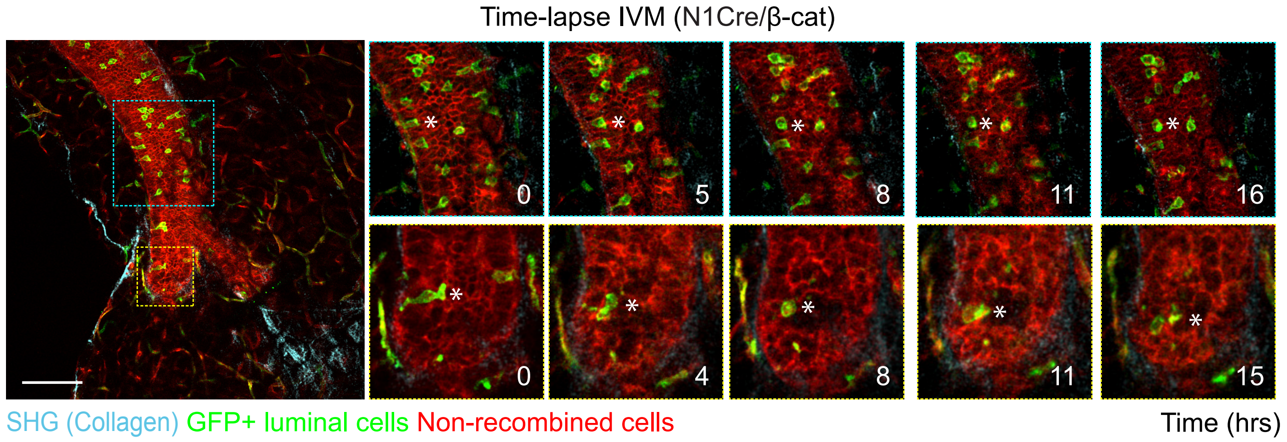


B

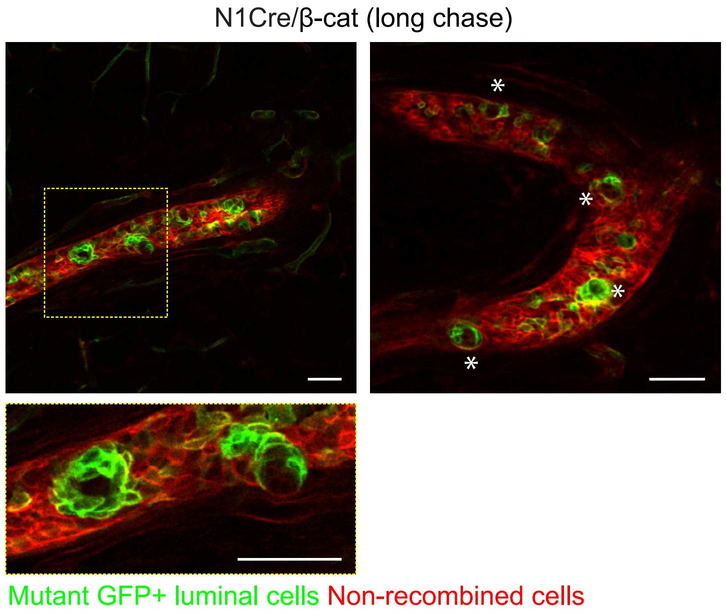

**Fig. S3. Time-lapse and longitudinal intravital imaging of mutant β-cat GFP+ luminal cells. Related to Fig.1.**

**(A)** Acute time-lapse IVM of luminal GFP+ (green) cell behaviors in a mammary duct of a pubertal N1Cre/β-cat mouse performed 48 h after low-dose tamoxifen administration. Red: non-recombined membrane tdTomato-expressing mammary epithelial cells; cyan: collagen (SHG). Related to Supplemental Movies 3 and 4. Asterisks denote cells being tracked over time. **(B)** IVM images (single *z*-planes) of luminal GFP+ lesions in a mammary duct of a N1Cre/β-cat mouse 19 (left panel) and 21 (right panel) days after low-dose tamoxifen administration. Asterisks demark loss of fluorescence in the center of ring-like lesions. Scale bars: 100μm in A and 50μm in B.

Supplementary Figure 4

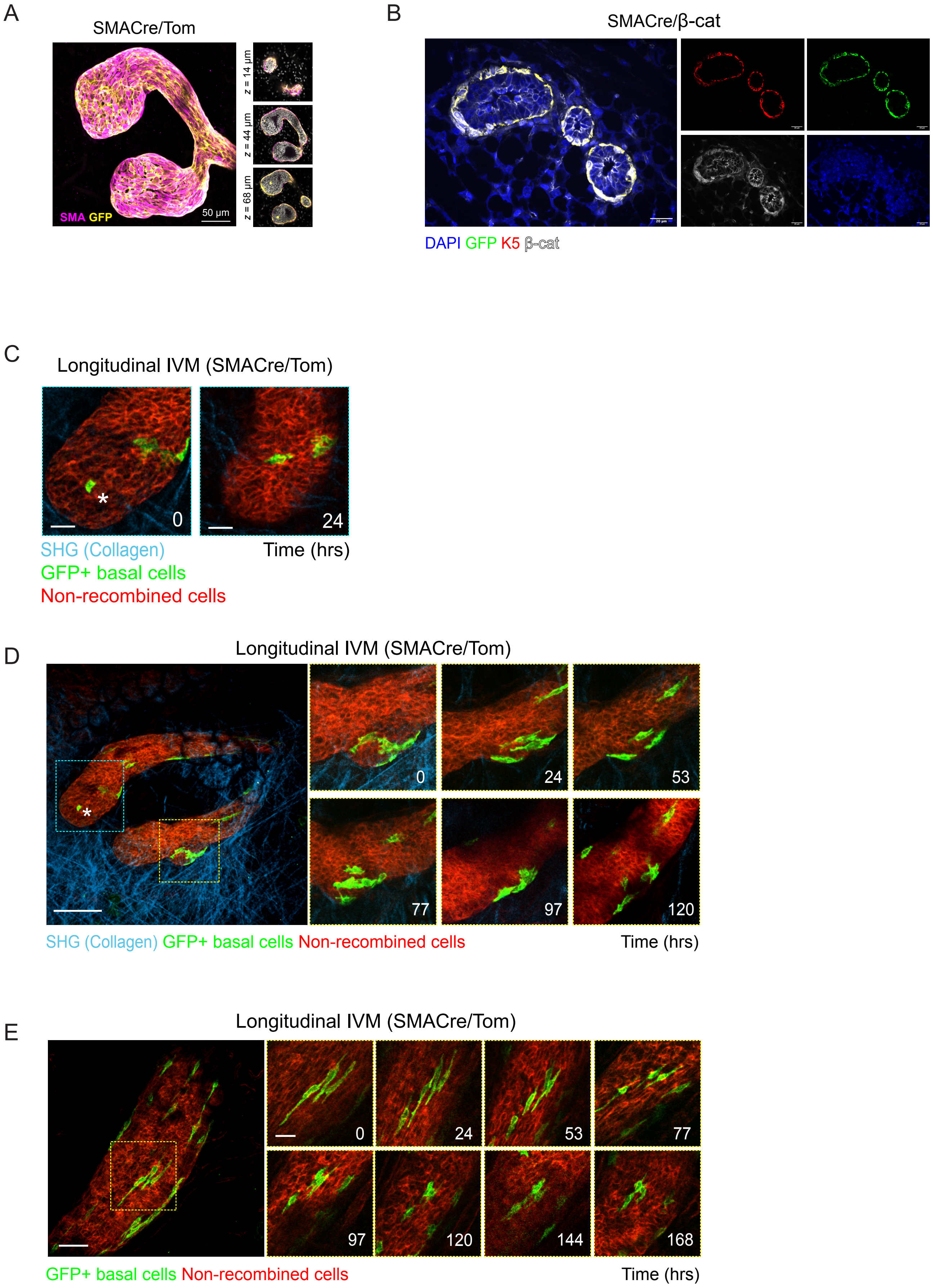

**Fig. S4. Longitudinal IVM of wildtype basal cell behaviors during pubertal ductal development. Related to Fig.2.**

**(A)** Maximum intensity z-projection and thin optical slices (depth in *z* relative to the first image in the image sequence) of cleared mammary tissue from SMACre/Tom mice immunostained with smooth muscle actin (SMA in pink). GFP+ cells (shown in yellow) are restricted to the basal compartment. **(B)** Representative sections of mammary tissues showing β-catenin accumulation in K5-expressing basal cells (in red) in SMACre/β-cat mice, coinciding with membrane GFP expression (in green). Anti-β-catenin staining is in white and DAPI labels nuclei in blue. **(C)** IVM images of a mammary end bud structure in SMACre/Tom mice showing the elimination of basal GFP+ (green) cap-in-body cells within 24 h. Inset of Fig. S4D (blue box). Red: non-recombined membrane tdTomato-expressing mammary epithelial cells; cyan: collagen (SHG). **(D)** IVM images of a mammary end bud structure in a pubertal SMACre/Tom mouse showing recombined GFP+ (green) mammary basal epithelial cell rearrangements over time (120 h). The asterisk denotes cap-in-body cell shown in Fig.S4C. Red: non-recombined membrane tdTomato-expressing mammary epithelial cells; cyan: collagen (SHG). **(E)** IVM images of ductal basal GFP+ (green) cells in the mammary gland of a pubertal SMACre/Tom mouse over time (168 h). Red: non-recombined membrane tdTomato-expressing mammary epithelial cells. Scale bars: 50μm in A, E; 20μm in B; 25μm in C and inset in E and 100μm in D.

Supplementary Figure 5

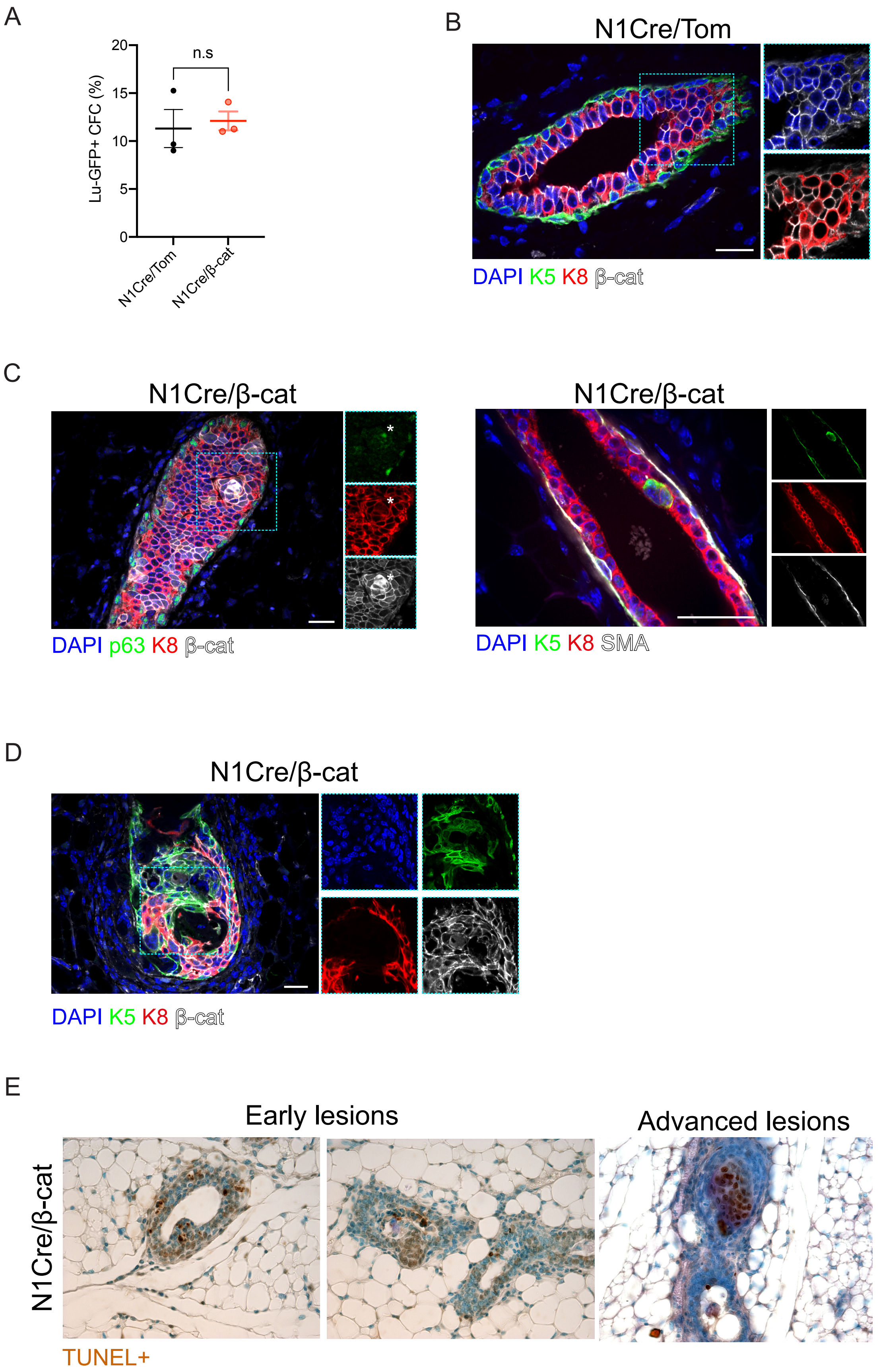

**Fig. S5 β-catenin stabilization in mammary luminal cells leads to the development of lesions with aberrant lineage marker expression and eventual cell death. Related to Fig. 3.**

**(A)** Quantification of colony forming assay showing no differences in the colony forming capacity of GFP+ luminal cells isolated by flow cytometry from the mammary glands of N1Cre/Tom and N1Cre/β-cat mice. n.s: not significant. **(B)** Representative sections of mammary ducts in N1Cre/Tom mice showing β-catenin expression (white) restricted to the cell membrane. **(C)** Representative sections of N1Cre/β-cat mammary ducts showing acquisition of p63 and K5 basal marker expression (but not SMA) in hyperplastic lesions. **(D)** Representative image of an advanced lesion in a N1Cre/β-cat the mammary duct showing the formation of rosette-like structures. **(E)** TUNEL staining in N1Cre/β-cat mammary sections indicating that cells within early and advanced hyperplastic lesions undergo cell death. Scale bars: 20 µm.

Supplementary Figure 6

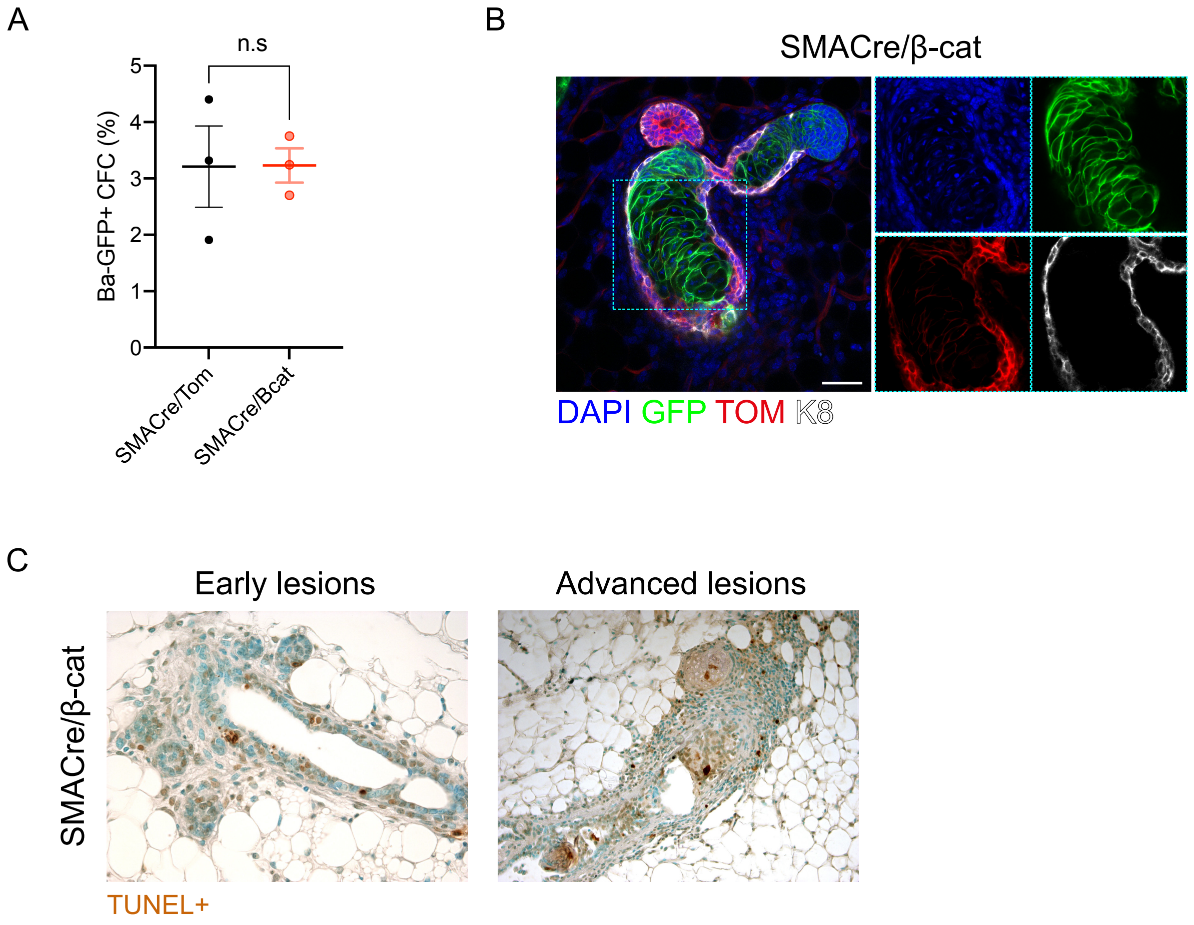

**Fig. S6 β-catenin stabilization in mammary basal cells leads to development of rosette-like lesions and eventual cell death. Related to Fig. 4.**

**(A)** Quantification of colony forming assay showing no statistically significant differences in the colony forming capacity of GFP+ basal cells isolated by flow cytometry from the mammary glands of SMACre/Tom and SMACre/β-cat mice. n.s: not significant. **(B)** Representative image of an advanced lesion in a SMACre/β-cat mammary duct showing formation of rosette-like structures with aberrant nuclei. Scale bar: 50 µm. **(C)** TUNEL staining in SMACre/β-cat mammary sections indicating that cells within early and advanced hyperplastic lesions undergo cell death.

Supplementary Figure 7

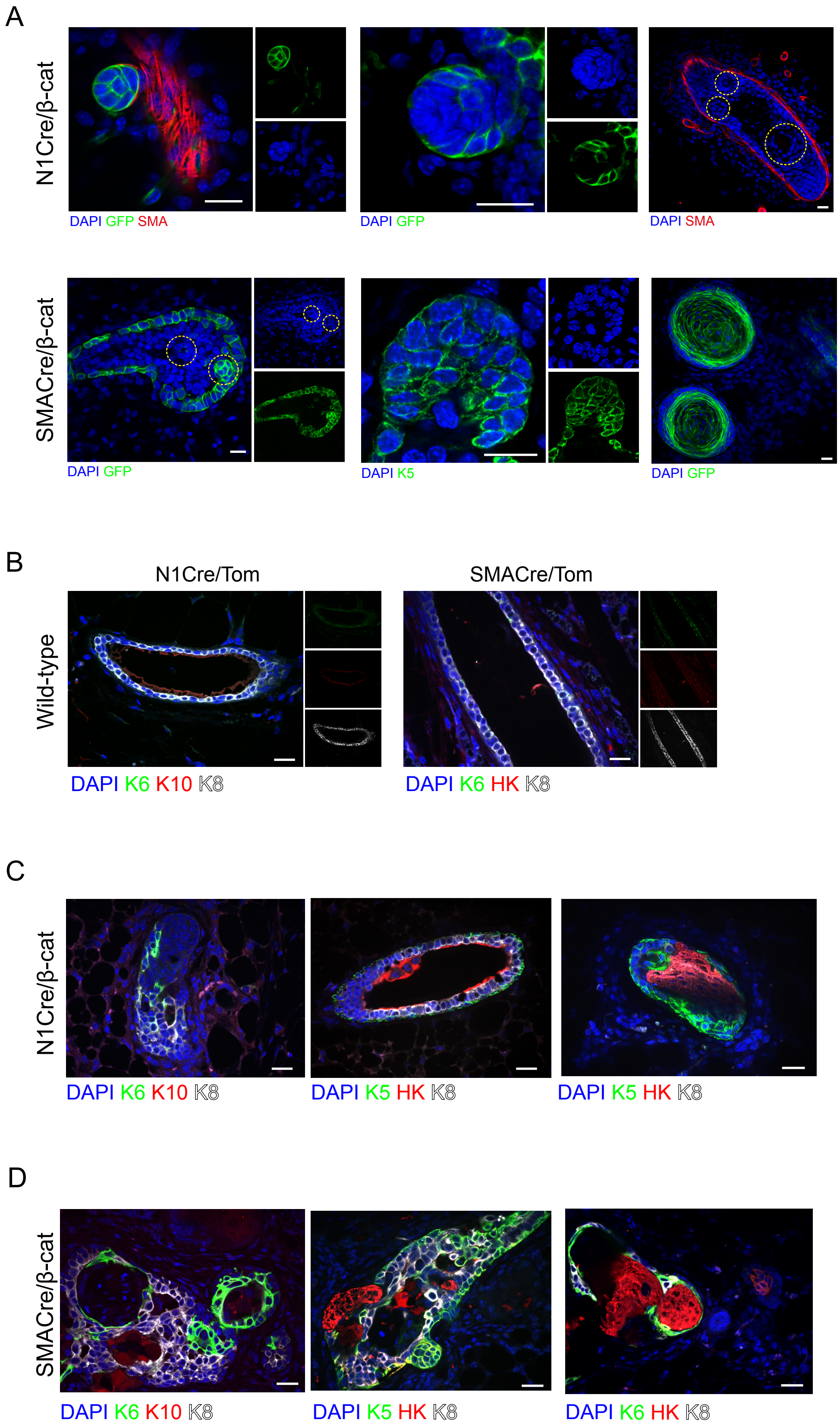

**Fig. S7 β-catenin stabilization drives the acquisition of a squamous transdifferentiation program in mammary epithelial cells, regardless of the targeted lineage.**

**(A)** Representative confocal images of β-catenin-expressing cells (in green) arranged into ring and bud-like clusters in early and advanced lesions in the mammary glands of N1Cre/β-cat and SMACre/β-cat mice. **(B)** Representative sections of wild-type mammary glands showing the lack of expression of the hair follicle markers K6, K10 and Hair Keratin (HK). **(C)** Representative sections of N1Cre/β-cat mammary glands showing acquired expression of the hair follicle markers K6 and Hair Keratin (HK) **(D)** Representative sections of SMACre/β-cat mammary glands showing expression of the hair follicle markers K6 and Hair Keratin (HK). Mammary epithelial cells in both models do not express the skin differentiation marker K10. Scale bars: 20 µm.

Supplementary Figure 8

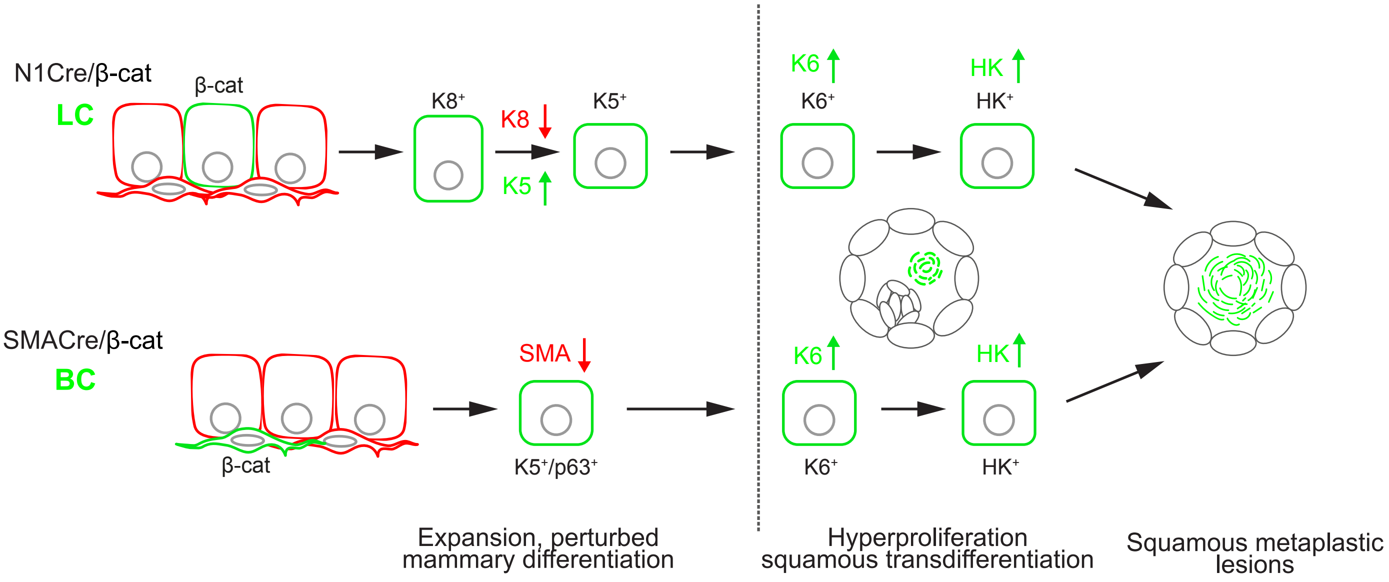

**Fig. S8.** Schematic representation of the timeline of squamous transdifferentiation induced by $\beta$-catenin stabilization in luminal (top) or basal (bottom) mammary epithelial cells.

**Supplemental Movie Legends**

**Supplemental Movie 1:** Longitudinal intravital imaging of a mammary end bud structure in a pubertal N1Cre/Tom mouse showing the dynamic cellular rearrangements and behavior of GFP+ luminal epithelial cells over 6 days (144 h). Red: non-recombined membrane tdTomato-expressing mammary epithelial cells; cyan: collagen (SHG). Related to Figure 1.

**Supplemental Movie 2:** Acute time-lapse IVM of GFP+ luminal epithelial cells residing in a mammary end bud structure of a pubertal N1Cre/Tom mouse. Red: non-recombined membrane tdTomato-expressing mammary epithelial cells; cyan: collagen (SHG). Total movie length is 06:00 (h:min). Related to Figure 1.

**Supplemental Movie 3:** Acute time-lapse IVM of GFP+ luminal epithelial cells in the mammary gland of a pubertal N1Cre/β-cat mouse. Red: non-recombined membrane tdTomato-expressing mammary epithelial cells. Total movie length is 16:30 (h:min). Related to Fig. 1 and S3.

**Supplemental Movie 4:** Acute time-lapse IVM of GFP+ luminal epithelial cells in the mammary gland of a pubertal N1Cre/β-cat mouse. Red: non-recombined membrane tdTomato-expressing mammary epithelial cells. Total movie length is 16:00 (h:min). Related to Fig. 1 and S3.

**Supplemental Movie 5:** Acute time-lapse IVM of GFP+ basal epithelial cells in the mammary gland of a pubertal SMACre/β-cat mouse. Red: non-recombined membrane tdTomato-expressing mammary epithelial cells. Total movie length is 12:30 (h:min). Related to Figure 2.

**Supplemental Movie 6:** Longitudinal intravital imaging of a mammary end bud structure in a pubertal SMACre/β-cat mouse revealing the expansion of mutant GFP+ cap-in-body epithelial cells over 9 days (216 h). Red: non-recombined membrane tdTomato-expressing mammary epithelial cells; cyan: collagen (SHG). Related to Figure 2.
